# Life and work of researchers trapped in the COVID-19 pandemic vicious cycle

**DOI:** 10.1101/2021.02.02.429476

**Authors:** S. Aryan Ghaffarizadeh, S. Arman Ghaffarizadeh, Amir H. Behbahani, Mohammad Mehdizadeh, Alison Olechowski

## Abstract

COVID-19 has disrupted researchers’ work and posed challenges to their life routines. We have surveyed 740 researchers of which 66% experienced a decrease in productivity, 50% indicated increased workload, and 66% reported they have been feeling internal pressure to make progress. Those whose research required physical presence in a lab or the field experienced considerable disruption and productivity decrease. About 82% of this group will try to permanently reduce their work dependency on physical presence. Parents and those taking care of vulnerable dependents have been spending less time on research due to their role conflict. We further observed a gender gap in the overall disruption consequences; more female researchers have been experiencing a reduction in productivity and external pressure to make progress. The results of this study can help institution leaders and policymakers better understand the pandemic’s challenges for the research community and motivate appropriate measures to instill long-term solutions.

## Introduction

The COVID-19 pandemic has affected the life and work of researchers on an unprecedented global scale. Early studies have shed light on the magnitude of the pandemic’s disruption on scientific progress by investigating the change of allocations of work time[1] and the means and effectiveness of communication during the remote working environment of researchers[2].

During this pandemic, researchers have developed new life and work routines based on constantly changing circumstances such as access to work-space, teaching duties, and parenting responsibilities, to name a few. Despite recent major progress in vaccine development and deployment, long-term fundamental shifts in the life and work routine of researchers such as the utilization of online platforms and reducing non-essential travels are already being predicted[3, 4]. Nevertheless, there has been a lack of research evidence collected regarding the evolving work-life balance routines of researchers in the past months.

This knowledge is crucial to policymakers as well as the research community to adjust current guidelines. We hope that by following the constructed guidelines, the scientific community can mitigate the long term negative effects and devise appropriate preventive measures for future global-scale crises.

We have conducted a survey to better understand the current life and work routine of researchers, as well as the long term effect of the pandemic on research. Within a 10 day period starting on December 1, 2020, 740 responses were collected from active researchers across North America and Europe.

## All researches were disrupted

Overall, 71% of participants’ institutions experienced a full closure-no physical access-for at least one month for non-essential personnel. Only 3.5% of participants had full access to their work space at all times, while the rest had partial access. At the time of data collection, the portion of researchers with no-access reduced to 16%, and the partial access was increased to 51%, Figure 1. A central stratifying measure in our study was whether physical presence in a lab or in the field is needed for the researcher to conduct their work. About 36% of participants indicated that the general state of their research requires physical presence in a lab or in the field, while the rest indicated having access to internet and computational resources are sufficient to conduct their research work. For the rest of this paper, the former group is going to be addressed as the Physical Presence Group (*PPG*) and the latter as the No Physical Presence Group (*NPPG*). Within the PPG, 94% indicated either minor or major disruption in their work by the closure period. Only 66% of NPPG’s population experienced such interruption, which is still noticeable for a group whose research nature does not require physical presence. The main factor for interruptions for NPPG, we reported to be lack of access to resources (computational, library, software, etc.).

**Figure 1:**
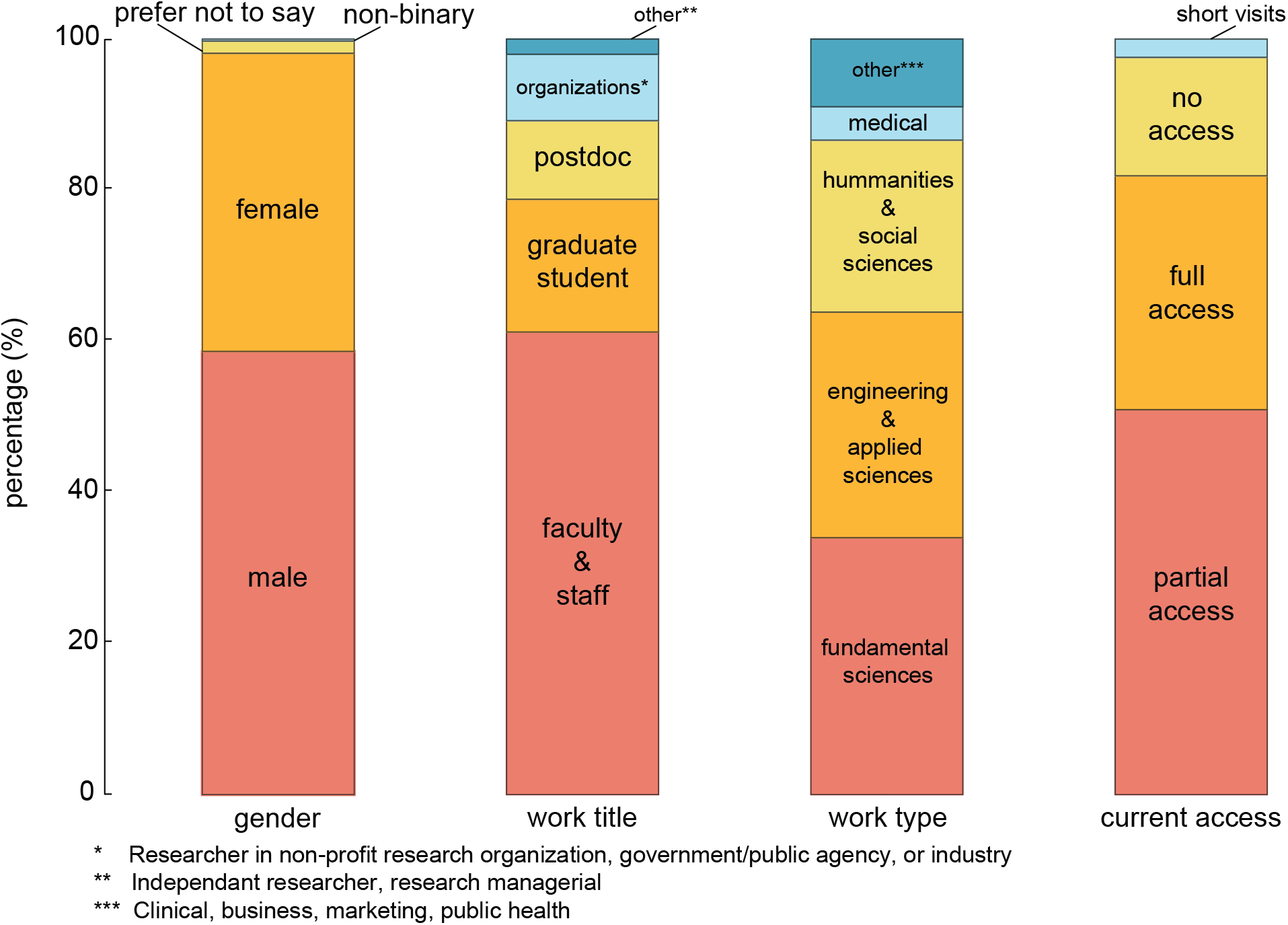
The percentage is based on 740 gathered responses; 639 participants from North America and 101 from Europe.

During the reopening stage, institutions have been prioritizing individuals and groups whose work have been impacted the most. These groups consist of experimental workers with perishable samples, time-sensitive data measurements, funding deadlines, etc. Our data reveals that a high degree of progress even for the NPPG depends on their PPG collaborators. Hence, institutions should consider the interconnections and dependencies among various projects and research groups in the next stages of reopening.

## Reduced Productivity: A general trend

The effect of local or global shocks and uncertainties on individual and societal level productivity has been studied in many contexts including wars, financial crisis, and natural disasters[5, 6]. The reallocation of resources from low to high productivity units at the times of uncertainty affects the productivity growth on large scales.[7] On an individual level, the shift of priorities and responsibilities can affect productivity. In our study, a general trend across all responses, when asked about the change of productivity was a significant reduction in productivity. About 66% of all participants reported a drop in productivity. With a higher overall average, 76% of the PPG experienced a drop while 60% of the NPPG believed that their productivity was reduced, Figure 2. The 16% difference between the two groups in lowered productivity may be associated with a smoother transition for the NPPG to a remote workplace. Also, across all groups, trouble adapting to the new lifestyle was the main source of productivity reduction, reported by 74% of respondents. Second to lifestyle adaptations, lack of motivation (with 51%) was another major contributor to decreased productivity. While the majority of our data show a reduction in productivity, 12% of the PPG and 16% of the NPPG experienced productivity increase. Within the NPPG and PPG with increased productivity, 56% and 39% of participants indicated not commuting to be the main reason for a higher productivity, respectively. In addition, for both groups, being more comfortable to work from home with 48% and 58% was the other notable reason for increased productivity, respectively.

**Figure 2:**
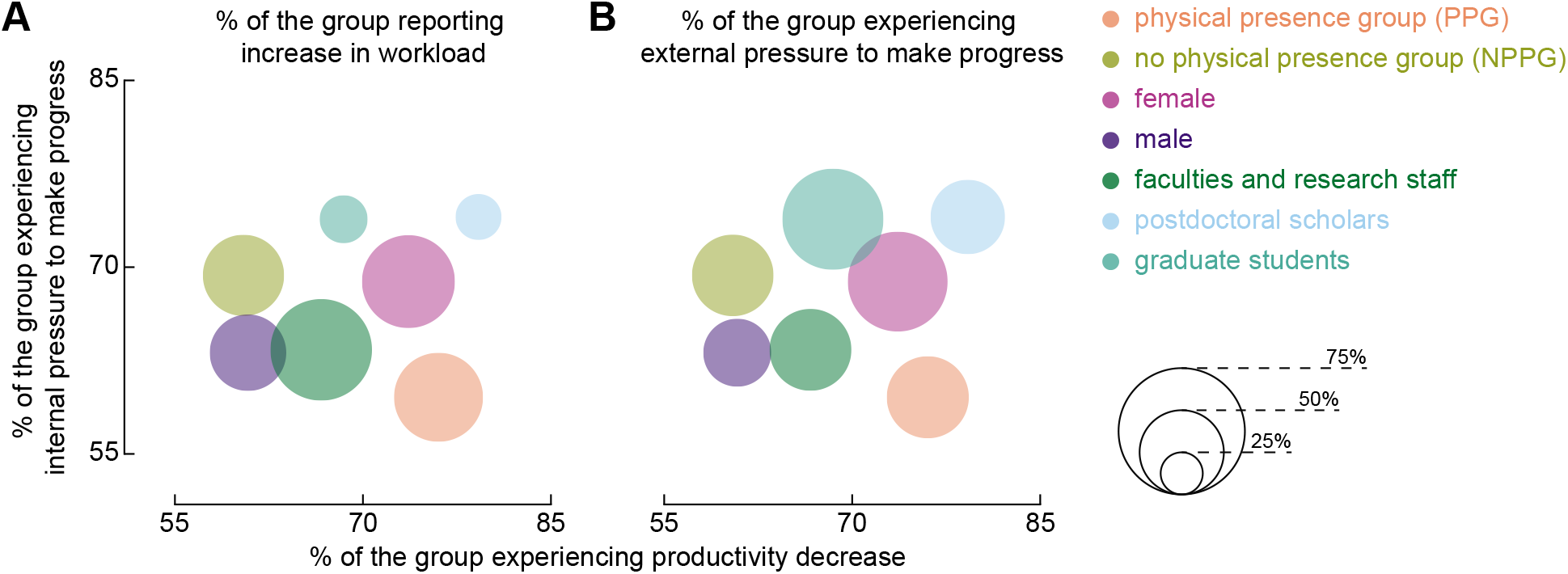
(A) The marker size indicate the percentage of each group experiencing workload. (B) The marker size indicates the percentage of each group experiencing external pressure to make progress.

Some of the practices that researchers got used to them during the pandemic can be continued even after the full return. For example, utilizing online platforms for holding meetings and events open up new opportunities, such as reaching to a broader audience[8].

## Female researchers were impacted more by non-work related responsibilities and external pressures

While the gender responsibility gap has become narrower over the past decades, recent studies showed that mothers compared to fathers spend 10 more hours per week on housework and childcare[9]. In addition to this workload, Offer and Schneider discuss that multitasking at work is associated with negative emotions and stress associated with feeling guilty about spending less time with family. Yavorsky *et al.* show that pre-birth data suggests an egalitarian division of housework, while post-birth, women spent significantly more time on childcare, housework, and total work[10].

A reasonable prediction is that the pandemic-driven closure of schools will put more burden on female researchers with parenting responsibilities[11]. Some parameters that could help us measure the hypothesis are productivity decrease, workload increase, and publication decrease. With the exception of reduced publications, which might surface only in the longer term due to the time frame of a publication process, the productivity and the workload can be measured. Evaluating the effect of demographic factors, female-identified participants reported a 74% productivity reduction while those identified as male experienced a 61% reduction. Analyzing the decreased productivity reasons, the gap among genders can be partially due to the higher number of females who were engaged in taking care of a child or a vulnerable person. Within female researchers, 27% have indicated that a combination of care-taking responsibilities has lowered their productivity, while 21% of male researchers were affected by these duties. Also, regardless of the productivity change, when asked about their new work-life routine, 26% of females indicated devoting more time to “taking care of my vulnerable dependant(s) (kids, parents, etc.)” in comparison to 16% of males.

We next studied the amount of internal and external pressures each group felt for progression in their work. For both females and males, the internal pressure to progress was 69% and 63%, respectively. This gap increases in the amount of external pressure each group felt. Approximately, 59% of female researchers indicated feeling external pressure to progress while 40% of males felt such a pressure, Figure 2.

Pandemic shed light on preexisting financial and mental challenges of simultaneously being a parent and a researcher. These challenges also reflected in career trajectory of female scientists seeking tenure track positions[12]. While such challenges are still widely unexplored, it is clear that the research community, especially graduate students, suffer from the lack of supportive policies regarding affordable childcare programs.[13, 14] Also, this pandemic showed how unreliable common care-taking practices, such as relying on family members’ help, might be in uncertain times. A university-wide subsidized childcare program is an absolute necessity for encouraging more female students to pursue academic careers.

## A scientific pause can bring new insights

Due to limited physical access to the workplace, the process of data collection was interrupted and many researchers have been revisiting previously gathered data. Hence, we asked whether and how the reexamination of old data changed previously drawn conclusions. About 52% of the researchers who needed physical presence in the lab or a field indicated revisiting old data. Among this group, 52% mentioned that the reexamination of the previously gathered data led to a new idea, insight, or conclusion. About 97% found the new idea, insight, or conclusion to be supporting or complimentary compared to the previous ones. Only one participant found the new conclusion to contradict previous ones.

## Faculties experienced a higher workload increase

While new online platforms were created and the existing ones have increased their features to facilitate a smooth transition from in person activities to remote, not all groups benefited equitably. Among graduate students and postdocs, 28% and 27% of them indicated that their workload have increased, respectively. However, 60% of faculties and research staff with teaching and administrative responsibilities reported increased in workload. Among the faculties with increased workload, 21% have been spending more time on professional services and reviewing publications. Also, 26% have been devoting more time to take care of my vulnerable dependant(s) (kids, parents, etc.). Such difference is a direct result of life-stage differences of faculty members and graduate student and postdocs.

## The pandemic will increase the demand of remote jobs

The study showed that the restrictions on physical travel and commuting to labs or the field have unequally affected researchers whose work depend on their physical presence. When the PPG were asked if upon returning to the lab/field, they would consider reducing the dependency of their work on physical presence, only 19% said that they will not. About 33% indicated that they will and can reduce their work dependency on physical presence, and the remaining said that they want to but cannot. This is an important result that demonstrates how the consequences of this pandemic have changed long term perspectives of researchers about the nature of their work and lifestyle. Finally, when asked about the researchers’ comfort regarding going back to the workplace with current precautionary measures, 63% said yes to their readiness of going back while the remaining 37% said no. Examining researchers’ level of comfort for going back to their workplace sheds light on the importance of the recent progresses on developing the vaccine and its central role to the continuation of research projects.

## Conclusion

The pandemic-driven closure of work spaces and schools impacted researchers on an unprecedented level. The majority of researchers we surveyed reported that they are trapped in the vicious cycle of reduced productivity, increased workload, and pressure felt to progress. To ensure a sustainable return to the pre-pandemic progress level, we believe that research institutes and policymakers should take many contributing factors such as the mental health of the employees and gender gap into account in addition to considerations on physical health.

## Author contributions

S.Aryan.G, S.Arman.G, AHB, and MM, designed the survey under the supervision of AO. S.Aryan.G analyzed and interpreted results. S.Aryan.G and S.Arman.G wrote the initial draft of the manuscript and all authors revised and edited the manuscript.

