## Supplementary material (methods) for "Life and work of researchers trapped in the COVID-19 pandemic vicious cycle"

### Supplementary Information

#### Life and work of researchers trapped in the COVID-19 pandemic vicious cycle reduced productivity, internal pressure to progress, and increased workload

S.Aryan Ghaffarizadeh [1] , S.Arman Ghaffarizadeh,[2] Amir H. Behbahani,[3] Mohammad Mehdizadeh,[4] and Alison Olechowski[5]

[1)] *Institute of Biomaterials and Biomedical Engineering, University of Toronto, Toronto, ON, Canada*

[2)] *Department of Mechanical Engineering, Carnegie Mellon University, Pittsburgh, PA, USA*

[3)] *Division of Biology and Biological Engineering, California Institute of Technology, Pasadena, CA, USA*

[4)] *Department of Mechanical Engineering, Louisiana State University, Baton Rouge, LA, USA*

[5)] *Department of Mechanical and Industrial Engineering, University of Toronto, Toronto, ON, Canada*

**Human participants ethics protocol**
The study protocol has been approved by the Institutional Review Board (IRB) from University of Toronto (#39786). Informed consent was obtained from all participants.

**Data availability**
No identifiable information including name, address, institution name, etc. was gathered from the participants. However, as data include sensitive and personal information, the IRB does not permit individual-level data to be shared either personally or publicly. For more information about the possibility of obtaining data, please contact University of Toronto IRB.

**Supplementary Method 1: Survey Sampling & Recruitment**
The goal of recruitment was to obtain a large and preferably random set of active researchers. Searching through Web of Science (WoS) data base provided by University of Toronto library for papers published from 2018 onward, we extracted 46,323 email addresses. Invitation emails were sent to all and of these, 41,017 were delivered and a number were bounced backed. 8,222 recipients opened the email, and from those 818 clicked through the link to access the survey. 613 completed the survey. Also, about 10 universities in Canada and US were contacted and asked to forward the invitation emails to their students. Total number of students who received invitations cannot be traced back, but 203 responses were collected. Since COVID-19 pandemic guidelines and extent of lockdown are different across countries, only responses North America and countries in Europe were selected for data analysis. This was to ensure consistency within the minimum COVID-19 measures across the responses. Filtering responses solely based on geographical location specified by the participants, 740 responses were used for the analysis.
We used Google Survey for the study. The detail of the survey can be found below. The follow up questions based on each answer is indicated by “Skip to question X”.

#### Supplementary Method 2: Additional Results

Here, we present the extended version of the results discussed in the survey. PPG refers to physical presence group, and NPPG refers to no physical presence group. If there is an "and" between two statements, it means a conditional counting of two groups. For example, productivity decrease, and PPG means the number of people who experienced decreased in productivity that are also within the PPG group. Alternatively, "or" means the summation of two values. To change such value to a percentage, they are divided by a value shown after the division sign. Total means 740 participants

| **Productivity** | |
|---|---|
| Productivity decrease of Total/ Total | 66.08% |
| Productivity Increase of Total/ Total | 15.00% |
| Productivity Not affected of Total/ Total | 18.92% |
| Productivity decrease and PPG / PPG | 76.03% |
| Productivity decrease and NPPG/ NPPG | 60.47% |
| Productivity Increased and PPG/ PPG | 12.36% |
| Productivity Increased and *NPPG*/ *NPPG* | 16.49% |

| **Productivity PPG** | |
|---|---|
| Productivity Increase and Not commuting and PPG/ PPG | 39.39% |
| Productivity Increase and Not commuting or more comfortable from home and PPG/ PPG | 87.88% |
| Productivity increase and more comfortable from home and PPG/ PPG | 48.48% |
| Productivity Increase and Not happy with workplace and PPG/ PPG | 3.03% |

| **Productivity NPPG** | |
|---|---|
| Productivity Increase and Not commuting/ NPPG | 56.41% |
| Productivity Increase and more comfortable from home/ NPPG | 57.69% |
| Productivity Increase and Not happy with workplace/ NPPG | 7.69% |

| **Workload (Total)** | |
|---|---|
| Workload Increase of Total/ Total | 49.86% |
| Workload decreased of Total/ Total | 9.32% |
| Workload stayed the same of Total/ Total | 40.81% |

| **Productivity Work Type** | |
|---|---|
| Productivity Decrease and Faculties/ Faculties | 66.67% |
| Productivity Decrease and Graduate student/ Graduate students | 68.46% |
| Productivity Decrease and Postdocs/ Postdocs | 79.22% |

| **Workload (PPG)** | |
|---|---|
| Workload Increase and PPG/ PPG | 52.81% |
| Workload decreased and PPG / PPG | 10.11% |
| Workload stayed the same and PPG/ PPG | 37.08% |

| **Workload (NPPG)** | |
|---|---|
| Workload Increase and NPPG / NPPG | 48.20% |
| Workload decreased and NPPG/NPPG | 8.88% |

| | |
|---|---|
| Workload stayed the same and NPPG/ NPPG | 42.92% |

| Workload (Male) | |
|---|---|
| Workload Increase of Total/ Total | 45.39% |
| Workload decreased of Total/ Total | 11.29% |
| Workload stayed the same of Total/ Total | 43.32% |

| Workload (Female) | |
|---|---|
| Workload Increase of Total/ Total | 55.14% |
| Workload decreased of Total/ Total | 6.51% |
| Workload stayed the same of Total/ Total | 38.36% |

| Workload (Worktype) | |
|---|---|
| Workload Increase and faculty/ faculty | 60.26% |
| Workload Increase and graduate student/ graduate student | 28.46% |
| Workload Increase and postdoc/ postdoc | 27.27% |

| Pressure (Total) | |
|---|---|
| Feeling internal pressure to progress of Total/ Total | 65.81% |
| Feeling external pressure to progress of Total/ Total | 48.51% |
| Not feeling pressure to progress of Total/ Total | 13.78% |

| Pressure (PPG) | |
|---|---|
| Feeling internal pressure to progress and PPG/ PGG | 59.55% |
| Feeling external pressure to progress and PPG/ PGG | 48.69% |
| Not feeling pressure to progress and PPG/ PGG | 5.62% |

| Pressure (NPPG) | |
|---|---|
| Feeling internal pressure to progress and NPPG/ NPGG | 69.34% |
| Feeling external pressure to progress and NPPG/ NPGG | 48.41% |
| Not feeling pressure to progress and NPPG/ NPGG | 18.39% |

| Pressure (Male) | |
|---|---|
| Feeling internal pressure to make progress and male/ male | 63.13% |
| Feeling external pressure to make progress and male/ male | 40.32% |
| Not feeling pressure to make progress and male/ male | 16.36% |

| Pressure (Female) | |
|---|---|
| Feeling internal pressure to progress and female/ female | 68.84% |
| Feeling external pressure to progress and female/ female | 59.25% |
| Not feeling pressure to progress and female/ female | 10.62% |

| Pressure and Worktype | |
|---|---|
| Feeling internal pressure to progress and Faculty/ Faculty | 63.36% |
| Feeling external pressure to progress and Faculty/ Faculty | 48.57% |

| | |
|---|---|
| Feeling internal pressure to progress and Postdoc/ Postdoc | 74.03% |
| Feeling external pressure to progress and Postdoc/ Postdoc | 44.16% |
| Feeling internal pressure to progress and Graduate student/ Graduate student | 73.85% |
| Feeling external pressure to progress and Graduate student/ Graduate student | 60.00% |

| Disrupted work | |
|---|---|
| Disrupted work and PPG/ PPG | 94.01% |
| Disrupted work and NPPG/ NPPG | 65.54% |

| Change Work type | |
|---|---|
| Want and will change work type and PPG/ PPG | 32.58% |
| Want but cannot change work type and PPG/ PPG | 48.69% |
| Will not change work type and PPG/ PPG | 18.73% |

| Previous Data | |
|---|---|
| Looking at previous data/ and PPG PPG | 51.69% |
| Looking at previous data AND new insight/ PPG and Looking at previous data | 52.17% |
| New insight AND supporting/ PPG and looking at previous data and new insight | 97.22% |

| Would you go back? | |
|---|---|
| Yes | 62.98% |
| No | 37.02% |
| Yes and feeling internal pressure to make progress/ Yes | 64.38% |
| Yes and feeling external pressure to make progress/ Yes | 45.28% |
| Yes and feeling no pressure to make progress/ Yes | 14.81% |

| Reasons for Productivity Decrease (gender) | |
|---|---|
| Female and productivity decrease and a child <6 or 6-18 or vulnerable/ female and productivity decreased | 26.51% |
| Male and productivity decrease and a child <6 or 6-18 or vulnerable/ Male and productivity decrease | 21.21% |
| Female and productivity decrease and lack of motivation/ female and productivity decreased | 52.56% |
| Male and decrease and lack of motivation/ Male and productivity decrease | 49.62% |
| Female and decrease and adapting to new lifestyle/ female and productivity decreased | 68.84% |
| Male and decrease and adapting to new lifestyle/ Male and productivity decrease | 77.65% |

| Work title and work type | |
|---|---|
| PPG and Faculty/Faculty | 32.45% |
| PPG and graduate students/ Graduate students | 44.62% |
| PPG and Postdocs/Postdoc | 44.16% |
